# A novel methodological framework for the assessment of the neural control of the shoulder using high-density surface electromyography

**DOI:** 10.1101/2025.09.15.676404

**Authors:** J. Greig Inglis, Silvia Rio, Hélio V. Cabral, Caterina Cosentino, Roberto Pagani, Tea Lulic-Kuryllo, Clark R Dickerson, Francesco Negro

**Affiliations:** Department of Clinical and Experimental Sciences, Università degli Studi di Brescia, Brescia, Italy; School of Physical Education and Sports, Universidade Federal do Rio de Janeiro, Rio de Janeiro, Brazil; Biomedical Engineering Program (COPPE), Universidade Federal do Rio de Janeiro, Rio de Janeiro, Brazil; Postgraduate Program in Rehabilitation Sciences, Universidade Federal do Rio de Janeiro, Rio de Janeiro, Brazil; Department of Mechanical and Industrial Engineering, Università degli Studi di Brescia, Brescia, Italy; Office of Research, Waterloo Regional Health Network, Kitchener, Canada; Department of Kinesiology and Health Sciences, University of Waterloo, Waterloo, Canada

**Author notes:** Corresponding author: Prof. Francesco Negro, Department of Clinical and Experimental Sciences, Università degli Studi di Brescia, Viale Europa 11, Brescia, 25121, Italy.

**Keywords:** motor unit discharge rate, HDsEMG spatial distribution, neural control, shoulder biomechanics

## Abstract

The complex function of the shoulder relies on the coordinated activation of small and large muscles, including the deltoid, pectoralis major, trapezius, and latissimus dorsi. However, detailed knowledge of their neuromuscular control remains limited. This study aimed to develop a methodological framework to investigate the neural control of the larger superficial shoulder muscles by combining a six-degree-of-freedom load cell attached to a robotic arm with high-density surface electromyograms (HDsEMG). Six healthy participants performed isometric contractions (abduction, adduction, flexion, and extension) at 30% of maximal voluntary contraction with the shoulder positioned at 30° and 65° of lateral abduction. HDsEMGs were recorded from the four muscles and analysed at the global activation, spatial distribution of activation and motor unit levels. Global activation was quantified using averaged normalized root-mean-square (RMS) amplitude and spatial distribution using coefficient of variation of the topographic maps. Moreover, HDsEMGs were decomposed into individual motor unit spike trains using convolutive blind source separation, and motor unit behaviour was characterized by mean discharge rate and spatial distribution of motor unit action potentials (MUAPs). RMS maps revealed action-specific activation within and between muscles, with the upper trapezius active across all tasks, while the anterior, middle, and posterior deltoid, clavicular pectoralis major, and latissimus dorsi were predominantly activated during abduction, flexion, and extension. Motor unit discharge rate also showed task-dependent activity. MUAP spatial distributions further showed distinct motor unit territories within arrays, suggesting region-specific recruitment strategies across actions. In conclusion, this framework demonstrates that individual motor unit activity can be reliably measured non-invasively in the superficial shoulder muscles. The approach provides a methodological basis for novel incorporation of neural control information into biomechanical models of shoulder function.

## 1.0 INTRODUCTION

Actions of the humerus require the complex activation of the muscles surrounding the glenohumeral joint to contract and, ultimately, cause the desired action. During these actions, the activation of large muscles such as the *trapezius, pectoralis major, latissimus dorsi,* and *deltoid* must be precisely coordinated with several smaller muscles, in particular the rotator cuff elements, to generate torque across multiple joints and adapt to changes in both task demands and musculoskeletal alterations, while also maintain glenohumeral stability in a delicate balance (Veeger & van der Helm, 2007). According to Hughes and colleagues (1999), the muscles of the glenohumeral joint (hereafter referred to as the shoulder) and their strength are important components of functional capacity. Thus, understanding the neuromuscular behaviour of these muscles is essential for a complete understanding of the activation and function of this muscle group, which can inform the prescription of appropriate training and rehabilitation strategies. Biomechanical modelling of the shoulder relies on simple mechanical analyses of body segments and the internal and external forces acting on the joint to predict shoulder muscle control (Hughes & An, 1997; Chang *et al*., 2000; Dickerson *et al*., 2008; Seth *et al*., 2019). However, to date, there are restrictions on modelling advances in the shoulder, particularly in including neural control of active muscles and, specifically, their partitional behaviour. Simplifying assumptions include single on/off activation of entire muscles or semi-arbitrarily defined mechanical elements. These limitations stem from the historical methodological approach to bipolar surface electromyogram recordings for inferring the intricate behaviour of the individual motor units involved in muscle activation. In fact, to date, the inclusion of motoneuron behaviour from the shoulder muscles in neuromechanical models remains elusive (Hughes & An, 1997). Therefore, shoulder models must rely on indirect measures of neural control, using metrics that only partially capture the complexity of muscle activation strategies (Farina *et al*., 2010; Lulic-Kuryllo *et al*., 2021*a*).

Classically, the approach to assessing muscular neural control has been global bipolar surface electromyography (EMG), using amplitude measures to infer neuromuscular adaptations to specific actions with applications in biomechanics (Lawrence & De Luca, 1983; Woods & Bigland-Ritchie, 1983; Hermans *et al*., 1999; Day & Hulliger, 2001; Karlsson & Gerdle, 2001; Keenan *et al*., 2005; Farina *et al*., 2010). However, the specificity of EMG amplitude in inferring neural drive to the muscle remains debated (Christie *et al*., 2009; Farina *et al*., 2010). Amplitude cancellation of surface motor unit action potentials is a key factor affecting the relationship between EMG amplitude and neural drive (Yao *et al*., 2000; Day & Hulliger, 2001; Dimitrova & Dimitrov, 2003; Keenan *et al*., 2005), because as additional motor units may be recruited during the gradation of force, the superposition of positive and negative phases of action potentials could result in a non-proportional increase in EMG amplitude (Day & Hulliger, 2001; Keenan *et al*., 2005). Additionally, bipolar EMG provides only muscle activity in the recording volume of the electrodes (i.e., local information). In the case of the shoulder, where large muscles produce complex actions across multiple degrees of freedom, not all muscle portions may be either monitored or active during all actions (i.e., within-muscle localized activation) (Lulic-Kuryllo *et al*., 2021*a*) via standard bipolar surface EMG. For instance, previous evidence suggested that localized regions of the deltoid, pectoralis major, and latissimus dorsi can be activated based on the specific shoulder action direction (Wickham & Brown, 1998, 2012; Brown *et al*., 2007; Lulic-Kuryllo *et al*., 2022). Finally, when recording myoelectric signals with global bipolar EMG, the risk of crosstalk must be considered. Signals generated by secondary muscles may lead to the erroneous interpretation of myoelectric signals from the primary muscle under investigation (De Luca & Merletti, 1988). Therefore, caution must be taken when assessing muscle activity using bipolar global signals, given the possibility of cancellation or crosstalk in the myoelectric recordings.

Advancements in myoelectric recordings in the early 1980’s have led to high-density surface electromyograms (HDsEMG) (Masuda *et al*., 1983, 1985; Reucher *et al*., 1987; Merletti *et al*., 1999). The benefit of recording with the evolving HDsEMG technology is its non-invasive nature and the ability to provide the spatial distribution of activation within and across muscles during complex actions. Additionally, when combined with motor unit decomposition algorithms (Holobar & Zazula, 2007; Ning *et al*., 2015; Chen & Zhou, 2016; Negro *et al*., 2016), the approach has the potential to record a large number of concurrently discharging motoneurons during voluntary contractions (Inglis *et al*., 2025). As the alpha motoneuron is the final common pathway of the neuromuscular system, integrating synaptic inputs from spinal and supraspinal pathways (Liddell & Sherrington, 1925; Sherrington, 1925), the analysis of motor unit discharge behaviour provides an effective way to assemble information on the generation and control of actions. A recent systematic review and meta-analysis revealed that most motor unit behaviour investigations have focused on the lower rather than the upper-body, leading to a paucity of data on the upper limb (Inglis *et al*., 2025). Additionally, it was revealed that of the upper limb muscles, specifically major movers of the shoulder, there are few motor unit behaviour studies investigating the *upper trapezius* (Westgaard & De Luca, 2001; Falla & Farina, 2008*a*, 2008*b*; Gerdle *et al*., 2008; Holobar *et al*., 2009; Dideriksen *et al*., 2016; Kirk *et al*., 2019) and *pectoralis major* (Lulic-Kuryllo *et al*., 2021*b*) using invasive or non-invasive techniques, none that examined the *latissimus dorsi* (Padovan *et al*., 2024) and only three examining the *deltoid* (De Luca *et al*., 1982; Doherty & Stashuk, 2003; Baslo *et al*., 2024), which all used intramuscular electrodes (Joseph *et al*., 2019).

Therefore, the aim of this paper was to develop a novel methodological framework to investigate four isometric shoulder actions (abduction, adduction, flexion and extension) at two shoulder elevation angles to fully assess and compare the neural control of the superficial shoulder muscles (deltoid, clavicular pectoralis major, upper trapezius and latissimus dorsi) at the global activation level (average HDsEMG amplitude), the spatial distribution of activation level (coefficient of variation of the HDsEMG amplitude maps) and the motor unit level (discharge patterns). For this, we combined a six-degree-of-freedom load cell with a robotic arm with the evolving HDsEMG technology. Additionally, this paper provides several examples of the type of extractable neural shoulder control information within the proposed framework. The novel framework proposed aims to establish guidelines for future investigations of the shoulder complex’s large muscles and to advance shoulder musculoskeletal modelling.

## 2.0 METHODS

### 2.1 Participants

Six healthy individuals free of any musculoskeletal or neuromuscular abnormalities (1 female; mean ± SD: age 28 ± 4 years; height 180 ± 7 cm; mass 79 ± 9 kg) voluntarily participated. All participants provided written informed consent before starting the experiments that conformed to the standards set by the latest version of the Declaration of Helsinki, which was approved by the ethics committee of the Università degli Studi di Brescia (code NP5890).

### 2.2 Experimental protocol

#### 2.2.1 Experimental setup

Each participant visited the lab on a single day where, in a randomized order, they performed multiple degree-of-freedom isometric contractions in a 2-hour session. Participants were seated in a rigid testing chair with their hip and knee joints at 90° of flexion (0° indicates full extension) and a broad Velcro® strap placed around the midsection to minimize lateral torso movements. The participant’s arm was extended and placed slightly (<20°) in the scapular plane to avoid discomfort when performing the selected actions. The participants’ upper arm was placed into a custom-built testing jig manufactured from rectangular metal tubing to ensure a rigid transfer of forces produced to the load cell (**Figure 1b**). The testing jig was attached to a 6-degrees of freedom load cell (© 2025 Advanced Mechanical Technology Inc., Watertown MA, USA, model #MC3A) to measure force output (**Figure 1c**). The load cell has a capacity of F_x_, F_y_ of 225 N and F_z_ of 445 N, and a sensitivity of F_x_, F_y_ of 5.4 µV/V-N and F_z_ of 1.35 µV/V-N. The load cell was attached to a robotic arm (Kawasaki Robotics Inc., Wixom MI, USA; **Figure 1a**) with a custom-machined aluminium coupling (**Figure 1d**). The importance of implementing the robotic arm was to position the testing jig and load cell through fine adjustments to accommodate individual anatomical landmarks, ensuring participants performed each action in identical relative positions and enabling precise recording of the forces produced during each action. Once positioned, the robotic arm was switched off and disconnected from all electrical sources to minimize electrical interference in the EMG signal.

**Figure 1.**
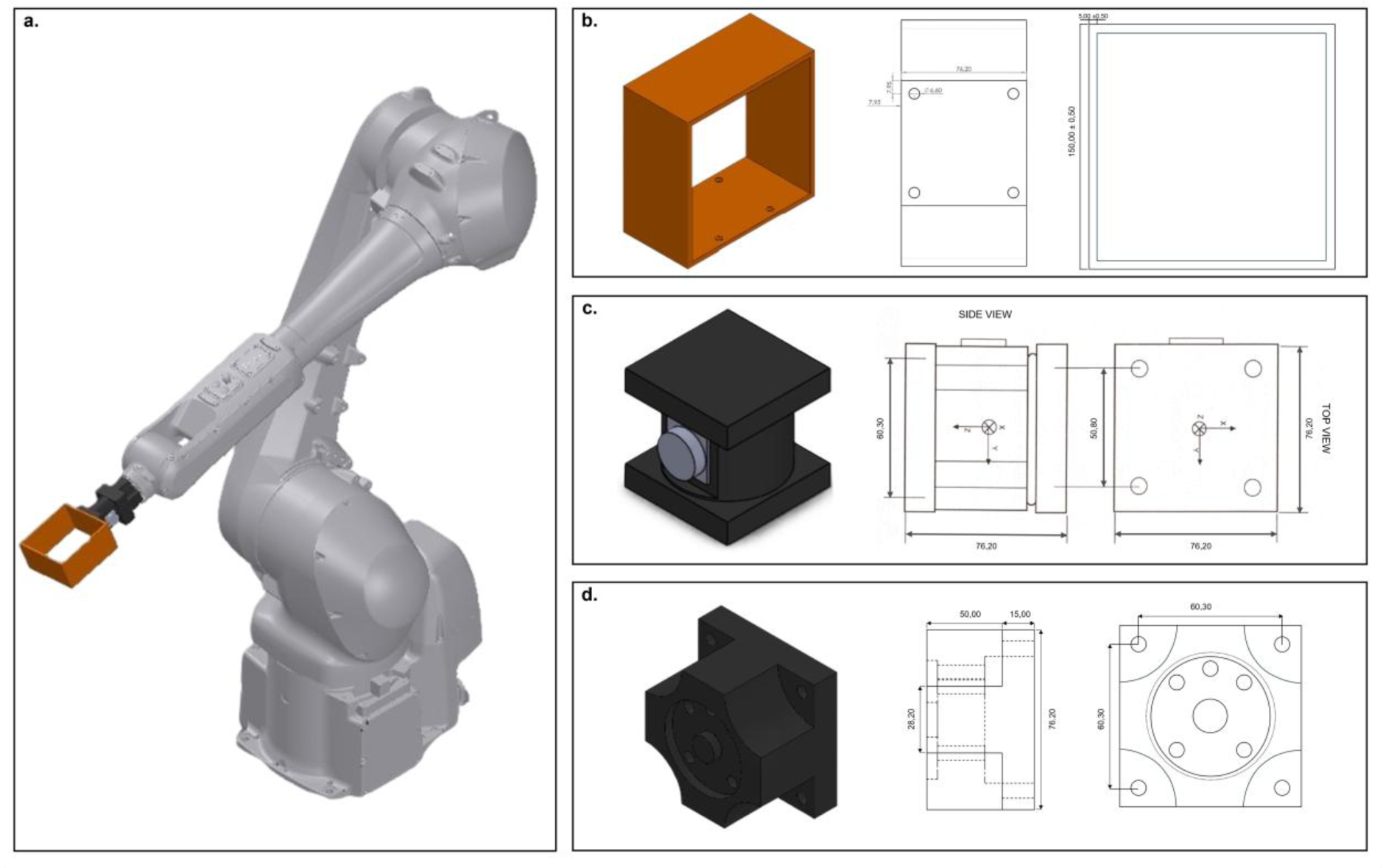
Mechanical components designed and manufactured for the robotic setup. (a) Complete robot assembly showing the robot with all components mounted. (b) 3D model and 2D technical drawings of the jig. (c) 3D model and 2D technical drawings of the six degree-of-freedom load cell. (d) 3D model and 2D technical drawings of the coupling between the robotic arm and the load cell. Measurements are reported in millimetres (mm).

#### 2.2.2 Experimental tasks

Each participant performed four isometric contractions at 30° and 65° of lateral shoulder abduction. The actions were: abduction, adduction, flexion and extension. **Figure 2** illustrates the position of the participant during the four actions across two shoulder angles. In **Figure 2**, the direction of the force vector is depicted with arrows for each action. Each action was performed at two shoulder angles to alter the deltoid muscle length (30° long; 65° short) and optimize the angles where the deltoid can initiate action during abduction (Peterson & Rayan, 2011).

**Figure 2.**
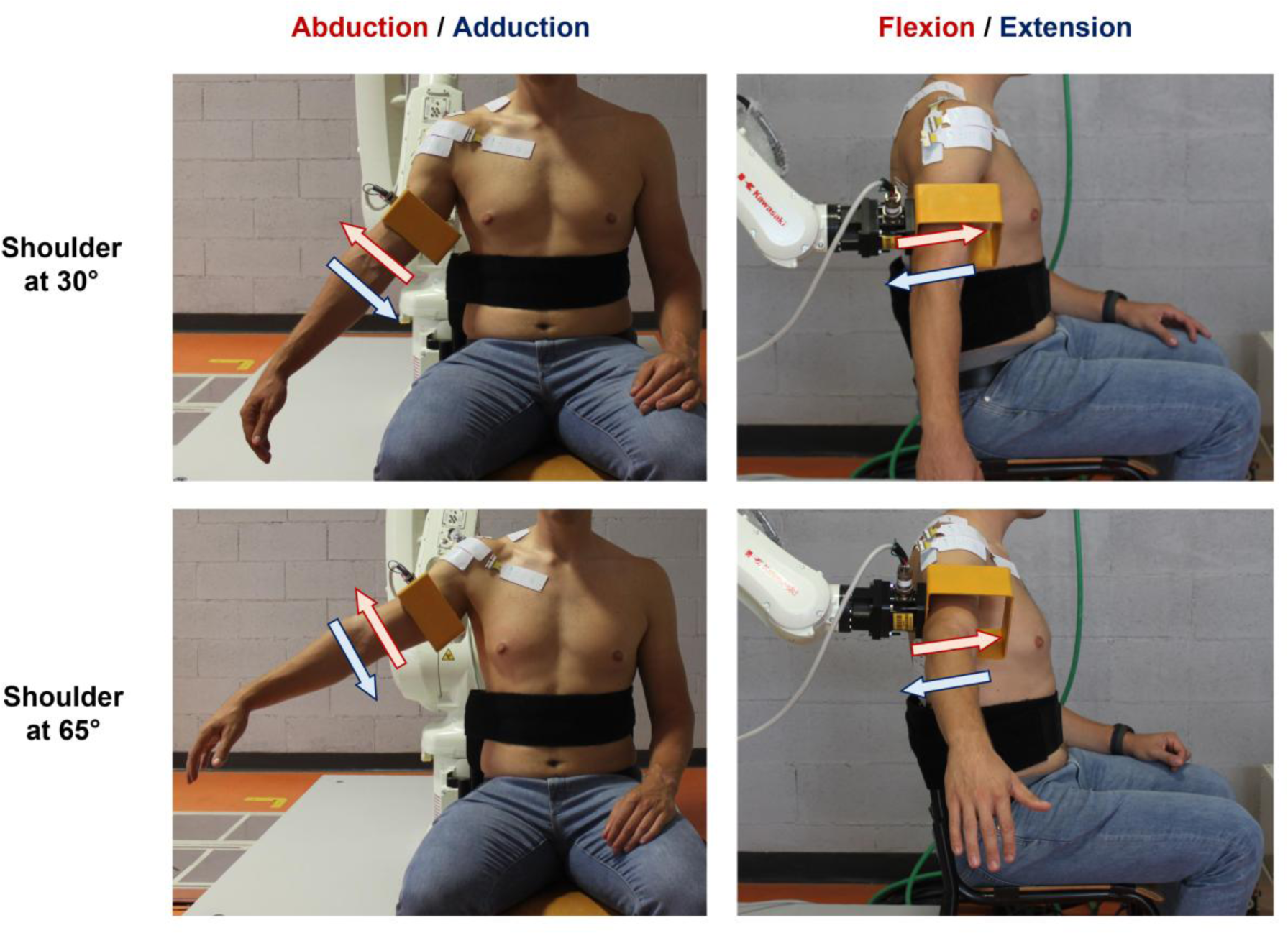
Experimental Actions. Shoulder abduction/adduction (left, anterior view) and flexion/extension (right, lateral view) performed at 30° (top) and 65° (bottom) of shoulder abduction. The arm was secured to a passive robotic interface, which secured the participant in position for each given action. A broad black Velcro® strap was placed around the midsection and secured to the testing chair to minimize actions of the trunk.

All experimental actions were repeated twice for each action and shoulder angle, resulting in a total of 16 trials (4 actions × 2 shoulder angles × 2 trials). Prior to each condition (angle/action), participants performed three voluntary maximum isometric contractions (MVCs) for 3-5 s, with a 3-min rest between each bout. The greatest value across the three MVCs was the maximal isometric shoulder force output for the individual condition and was used as a reference to set the target for the submaximal contractions.

Subsequently, participants underwent a short familiarization period, during which they were instructed to exert isometric shoulder forces at 10% MVC for approximately 10 s. This familiarization task was repeated up to a maximum of five times to ensure that participants could maintain the force output as steadily as possible at a target level, which was visually assessed by a researcher (JGI). Following the familiarization period, participants were instructed to perform two isometric contractions following a trapezoidal profile. The trapezoidal profile involved a linear increase at a rate of 3% MVC/s, a plateau phase for 30 s, and a linear decrease at a rate of 3% MVC/s. The target force output was set at 30% MVC (i.e., moderate force output) to mimic common outputs during activities of daily living (Lulic-Kuryllo *et al*., 2021*a*).

#### 2.2.3 Force feedback

A novel graphical user interface (GUI) was developed in MATLAB (version 2024a) to provide visual feedback of the force output and the rate at which the force was to be developed during the trapezoidal profile (**Figure 3e**). Force feedback was provided based on the axis being investigated relative to the three-axis load cell connected to the PC by an A-D board (National Instruments CORP., NI USB-6343). The selected axis was represented by either a horizontal or vertical line, where the Y direction was utilized for abduction and adduction, and the Z direction during flexion and extension (frontal) relative to the load cell. The GUI provided a green target box with the dimensions set to ± 2.5% MVC and a blue circle to represent the changes in force output. Each participant was instructed to maintain the blue circle inside the square. In cases where the error in force production exceeded 2.5% MVC, the square would turn red, and the participant would be encouraged to adjust the force output to meet the target force. Visual feedback was displayed on a monitor placed 1.3 meters in front of the participant at eye level.

**Figure 3.**
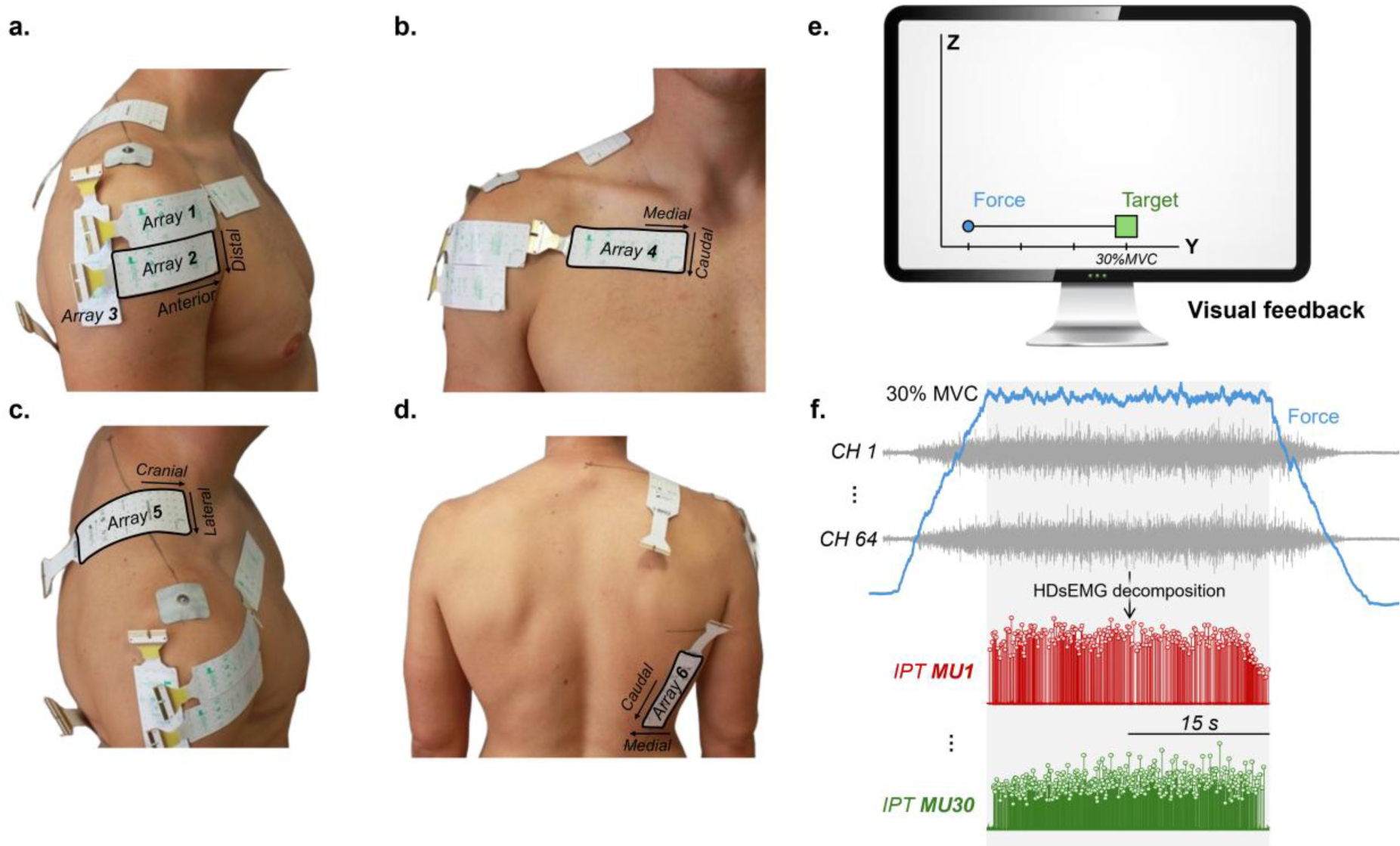
Electrode placement, visual feedback and HDsEMG analysis. (a) Electrode placement for the deltoid. Array 1 was positioned horizontally approximately 2 cm distal from the insertion of the deltoid tendon. Array 2 was placed distally and slightly offset to cover the lateral and anterior deltoid. Array 3 was positioned vertically adjacent to the distal deltoid array. (b) Electrode placement for the clavicular region of the pectoralis major. Array 4 was positioned horizontally and approximately 2 cm inferior to the clavicle, avoiding the sternum. (c) Electrode placement of the upper trapezius. Array 5 was positioned to bisect the 7^th^ and 8^th^ row of the array along the line between C7 and the acromion process. (d) Electrode placement of the latissimus dorsi. Array 6 was positioned diagonally in an area delimited by two horizontal lines, drawn approximately 3 cm inferior to the posterior axillary crease and parallel to the superior aspect of the iliac crest, the array was placed in line with the muscle fibres. (e) Visual feedback of the force output. Visual feedback was provided from a line, a square box and a circle representing the profile, the target force and the force output of the participant, respectively. (f) Schematic representation of the HDsEMG decomposition. HDsEMGs were visually inspected and, subsequently, decomposed into motor unit innervation pulse train (IPTs). The 30-s plateau phase was considered for this analysis (light grey rectangle).

### 2.3 Data collection

HDsEMG were acquired from the four major humeral movers (anterior, lateral and posterior deltoid, clavicular pectoralis major, upper trapezius, and latissimus dorsi) during the submaximal isometric actions using six two-dimensional adhesive arrays (HD08MM1305 and HD08MM1606; 1 mm diameter; 8 mm inter-electrode distance; OT Bioelettronica, Turin, Italy). The arrays placed over the deltoid and clavicular pectoralis major (see below) consisted of 64 electrodes arranged in 13 rows by 5 columns, with one missing channel in the top left corner. As for the upper trapezius and the latissimus dorsi, the arrays consisted of 96 electrodes arranged in 16 rows by 6 columns. Prior to array and reference electrode placement, the area was shaved, mildly abraded (EVERI, Spes Medica, Genova, Italy), cleaned with distilled water, and dried with paper towel. The electrode-skin contact was ensured by filling the foam cavities with conductive paste (AC cream, Spes Medica, Genova, Italy). Two reference electrodes were used; one positioned on the right elbow (electrode strap) at the level of the epicondyles and the other on the acromion process (snap electrode).

Several arrays and configurations were piloted to achieve the greatest motor unit yield in the deltoid (see appendix A). For the anterior/lateral deltoid, the electrode was placed approximately 2 cm distal to the proximal deltoid tendon, starting from the anterior border horizontally towards the posterior portion perpendicular to the insertion of the lateral fibres to the humerus (*array 1*, **Figure 3a**). To cover more of the anterior and lateral aspects of the deltoid, a second array was placed distally and slightly offset (2 array columns) and parallel to the proximal array (*array 2*, **Figure 3a**). For the posterior deltoid, the electrode array was positioned in line with the posterior aspect of the deltoid adjacent to the distal deltoid array (*array 3*, **Figure 3a**). For the clavicular region of the pectoralis major, the electrode array was placed approximately 2 cm inferior to the clavicle (superior *array* in (Lulic-Kuryllo *et al*., 2021*a*); *array 4*, **Figure 3b**). For the upper trapezius, the electrode array was positioned to bisect the 7^th^ and 8^th^ row (*array 5*, **Figure 3c**) along a line drawn between C7 and the lateral edge of the acromion. This was done because of the use of the 96-electrode array which is larger (3 extra rows and 1 extra column) than the standard 64 electrode array used for the deltoid and clavicular pectoralis major above (modified from (Falla & Farina, 2008*b*). For the latissimus dorsi, a line was drawn approximately 3 cm inferior to and horizontal to the posterior axillary crease and parallel to the superior aspect of the iliac crest. From these lines, a diagonal line was drawn to follow the latissimus dorsi muscle fibres, and the array was placed in the middle of this area, ensuring that the majority of the array covered the bulk of the muscle (Padovan *et al*., 2024) (*array 6*, **Figure 3d**).

HDsEMG and force outputs were digitized synchronously at a sampling frequency of 2000 Hz using a 16-bit A/D converter (10-500 Hz bandwidth; Novecento+, OT Bioelettronica, Turin, Italy). HDsEMG signals were recorded in monopolar derivation and amplified (8 V/V) to maximize signal resolution while avoiding saturation.

### 2.4 Data analysis

HDsEMG recordings were analysed offline in MATLAB (version 2024a) using custom-written scripts. As potential applications of the developed novel experimental framework, analyses were performed at the global activation level (average HDsEMG amplitude), the spatial distribution of activation level (coefficient of variation of the HDsEMG amplitude maps) and the motor unit level (discharge patterns) to assess the neural control of shoulder muscles. For all analyses, the 30-s plateau phase of each contraction was evaluated. HDsEMG were bandpass filtered between 20 and 900 Hz using a third-order Butterworth filter and subsequently visually inspected to identify and remove bad channels due to electrode-skin contact issues or artifacts (mean ± SD: 2 ± 1 channels removed across all actions and shoulder angles).

#### 2.4.1 HDsEMG root-mean-square amplitude and coefficient of variation

To assess the global activation of the investigated muscles, the root-mean-square (RMS) amplitude was utilized. Initially, single-differential (bipolar) EMGs were calculated as the algebraic difference between monopolar signals detected by consecutive rows of electrodes. Then, the RMS amplitude was calculated for each of the 59 (64-electrodes array) and 80 (96-electrodes array) single-differential channels over the duration of the trial, generating topographic maps of activation. The RMS amplitude maps were normalized to the average RMS value obtained from a 250 ms window, centred around the peak maximum isometric voluntary contraction, which was identified by a trained researcher (SR). RMS normalization was repeated for each muscle, action, and shoulder angle. For simplicity, only *array 2* of the deltoid was analysed. The average normalized RMS amplitude across all channels was then considered as the global RMS amplitude of the muscle for each action and shoulder angle.

To assess the variability in spatial distribution of the muscle activation, the coefficient of variation (CoV) of the normalised RMS amplitude maps was calculated (i.e., the standard deviation of the normalized RMS amplitudes divided by their mean). The CoV was calculated separately for each muscle, movement, and shoulder angle.

#### 2.4.2 HDsEMG decomposition

HDsEMG signals were completely decomposed into their constituent motor unit spike trains using a convolutive blind-source separation algorithm (Negro *et al*., 2016). Briefly, after extending and whitening the HDsEMG myoelectric signals, a fixed-point algorithm that seeks sources that maximize a measure of sparsity was applied to identify the motor unit spike trains (**Figure 3f**). The spikes were separated from the noise using the K-means algorithm and, while iteratively updating the motor unit separation vectors, the discharge times estimation was further refined by minimizing the coefficient of variation of the inter-spike intervals. This decomposition method has been previously validated (Negro *et al*., 2016) and extensively applied to assess the activity of single motor units (Negro *et al*., 2016; Cogliati *et al*., 2020; Cabral *et al*., 2024). After the automatic identification of motor units, any missed or misidentified motor unit discharges that produced non-physiological discharge rates (< 4 pulses per second or > 50 pulses per second; Negro *et al*., 2009) were manually identified and iteratively edited by an experienced operator (JGI, SR, CC), and the subsequent motor unit spike trains were re-estimated (Martinez-Valdes *et al*., 2017; Hassan *et al*., 2020). This approach has been shown to be highly reliable across operators (Hug *et al*., 2021). Only motor units with a silhouette (SIL) value, which is a quality control measure of the decomposition accuracy, greater than 0.87 were included in the further analysis (Negro *et al*., 2016).

#### 2.4.3 Motor unit discharge behaviour

The instantaneous discharge rates of each identified motor unit were determined by computing the inverse of the inter-spike intervals, and subsequently, the mean discharge rate was calculated and considered for further analysis.

#### 2.4.4 Motor unit action potential amplitude

To assess the spatial distribution of motor unit action potentials (MUAP), templates were obtained using the spike-triggered averaging technique (Stein *et al*., 1972). Specifically, single-differential EMGs were segmented into 100-ms windows centred on the discharge times of individual motor units and averaged. The peak-to-peak amplitude of each template was then calculated across channels, generating topographical MUAP amplitude maps. These maps were used both to validate the decomposed motor units and to exemplify the type of neural information that can be extracted with the proposed framework. As for the HDsEMG normalized RMS amplitude analysis, only *array 2* of the deltoid was analysed.

### 2.5 Statistical analysis

All statistical analyses were performed in R (version 4.2.2), using the RStudio environment. Linear mixed-effect models (LMM) were applied for all statistical analyses, as they account for the non-independence of data points within each participant, which is particularly relevant for motor unit data. To compare global normalized RMS amplitude, CoV of RMS amplitude maps and motor unit mean discharge rate, random intercept models were applied with action (abduction, adduction, flexion and extension) and muscle (deltoid, clavicular pectoralis major, upper trapezius and latissimus dorsi) as the fixed effects and participant as the random effect (e.g., *mean discharge rate ∼ 1 + action*muscle + (1 | participant)*). LMMs were implemented using the package *lmerTest* (Kuznetsova *et al*., 2017) with the Kenward-Roger method to approximate the degrees of freedom and estimate the *p*-values. The *emmeans* package was used, when necessary, for multiple comparisons and to determine estimated marginal means with 95% confidence intervals (Lenth *et al*., 2019). All individual data of motor unit discharge times are available at https://doi.org/10.6084/m9.figshare.30999811.

## 3.0 RESULTS

### 3.1 HDsEMG root-mean-square amplitude and coefficient of variation

**Figure 4a** depicts the topographical maps of the RMS amplitude at the 30° shoulder angle across muscle and actions for a selected participant. During abduction, the deltoid was the most active muscle, particularly in the lateral-anterior portion of *array 2* (see **Figure 3a**), followed by the caudal region of the upper trapezius. As for the clavicular pectoralis major and latissimus dorsi, during abduction, there was no visual evidence of significant activation. During adduction, most of the activity was in the latissimus dorsi and covered much of the array, followed by a small amount of activity in the medial-caudal portion of the clavicular pectoralis major array. However, there was no evident activity in the deltoid and upper trapezius during the adduction action. During flexion, the activity was predominantly in the anterior deltoid, followed by the clavicular pectoralis major. As for extension, the deltoid was again the most active, with the activity primarily towards the posterior aspect of the array. There was also significant activity in the caudal region of the array in the latissimus dorsi and upper trapezius, with no evidence of activity in the clavicular pectoralis major. In **Figure 4b**, the topographical RMS amplitude maps for the same selected participant are depicted, but at a shoulder angle of 65°. The activation patterns during abduction, adduction, and flexion closely resembled those reported above for the 30° condition. However, during extension, the deltoid was significantly active across *array 2*. Additionally, as seen for the 30° condition, a significant amount of activity was observed in the caudal region of the upper trapezius, but with little activity in the latissimus dorsi.

**Figure 4.**
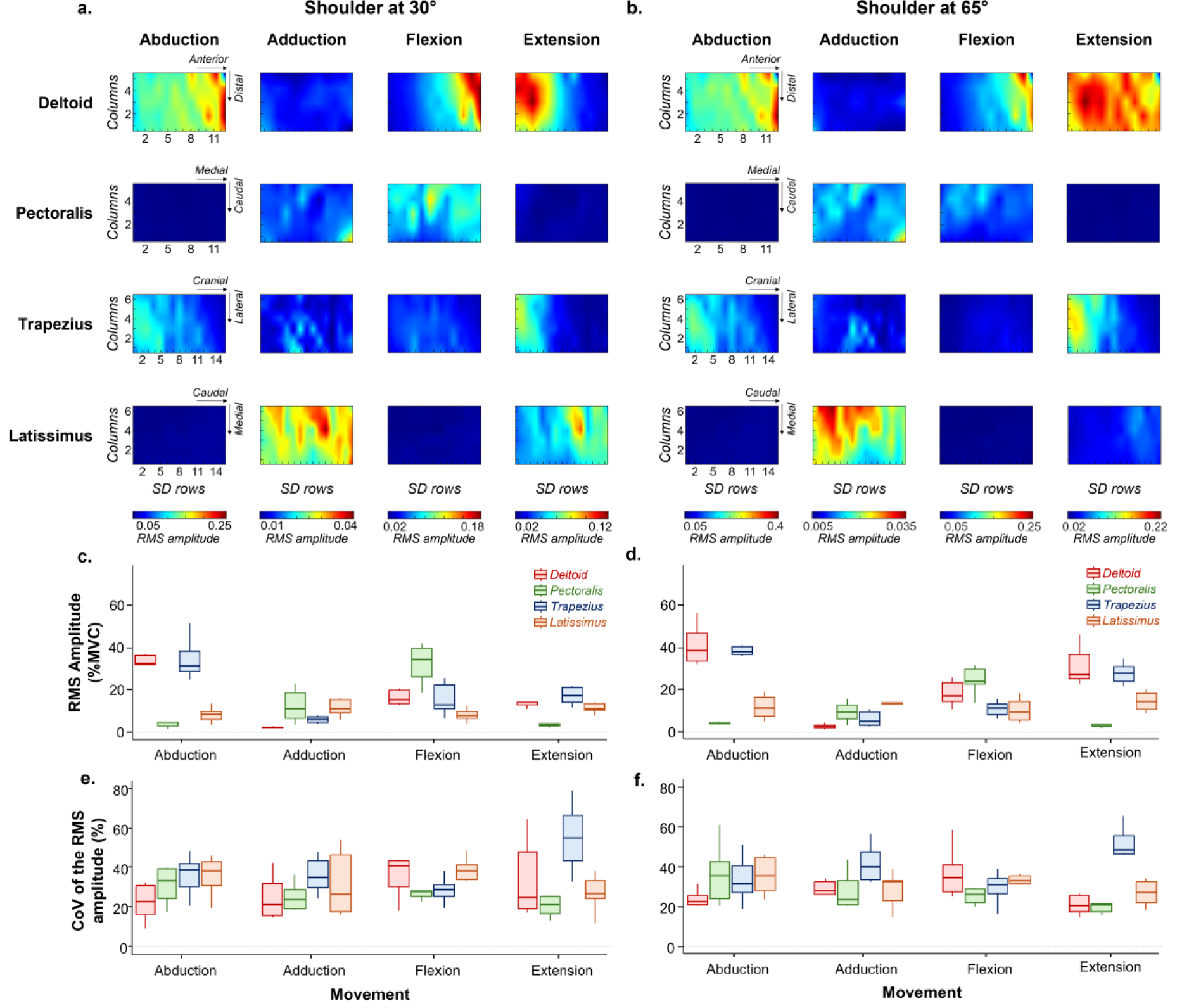
Muscle activation patterns across shoulder actions at 30° and 65°. RMS amplitude topographical maps for the deltoid, clavicular pectoralis major, upper trapezius, and latissimus dorsi during abduction, adduction, flexion and extension. These actions were performed with the shoulder positioned at 30° (a) and 65° (b) of lateral abduction. The x- and y-axes represent the rows and columns of the array, with colour intensity (from dark blue to dark red) indicating the level of muscle excitation. Panels (c) and (d) show the group results of the global normalized RMS amplitude (%MVC) for each action and muscle. Panels (e) and (f) show the group result of the coefficient of variation (CoV) of the RMS amplitude maps. Horizontal traces, boxes, and whiskers represent the median value, interquartile interval, and distribution range, respectively.

**Figures 4c** and **4d** show the group results for the global normalized RMS amplitude at 30° and 65° shoulder angles. For both 30° and 65° there were statistically significant interactions between muscle and action (30°: LMM, *F*=20.030, *p*<0.001; 65°: LMM, *F*=28.213, *p*<0.001). The following pairwise comparisons were common for both angles. For abduction, the deltoid and the upper trapezius had a significantly greater global normalized RMS amplitude compared to clavicular pectoralis major and latissimus dorsi (*p*<0.001 for all comparisons). During flexion, the normalized RMS amplitude of the clavicular pectoralis major was significantly greater compared to upper trapezius and latissimus dorsi (*p*<0.001 for all comparisons). Additionally, at the 30° shoulder angle, during flexion the normalized RMS amplitude of the clavicular pectoralis was significantly greater than the deltoid (*p*<0.001). During extension, both the deltoid (*p*=0.022) and the upper trapezius (*p*<0.001) had a greater normalized RMS amplitude compared to the clavicular pectoralis major, with no significant differences compared to the latissimus. When comparing the same muscle across actions at the 30° and 65° shoulder angle, the deltoid normalized RMS amplitude was significantly greater in abduction, flexion, and extension compared to adduction (*p*<0.01 for all comparisons), as well as during abduction compared to flexion and extension (*p*<0.01 for both comparisons). During flexion the clavicular pectoralis major exhibited the greatest normalized RMS amplitude compared to all other actions (*p*<0.001 for all comparisons). As for the upper trapezius, the normalized RMS amplitude was significantly greater during abduction compared to adduction, flexion, and extension (*p*<0.001 for all comparisons). Finally, the latissimus dorsi was not significantly different across all actions (*p*>0.05 for all comparisons).

The group results for the CoV of the normalized RMS amplitude maps at the 30° and 65° shoulder angle showed a statistically significant interaction between muscle and action (**Figure 4e and 4f**, 30°: LMM, *F*=2.790, *p*=0.007; 65°: LMM, *F*=2.789, *p*<0.001). The following pairwise comparisons were common for both angles. For abduction, adduction, and flexion there were no significant differences observed across muscle (*p*>0.05). During extension, the CoV of the normalized RMS amplitude maps of the upper trapezius was significantly greater compared to all other muscles (*p*<0.05 for all comparisons). While comparing the same muscle across movement, at the 30° shoulder angle, the upper trapezius had a significantly greater CoV of the normalized RMS amplitude maps during extension compared to all other movements (*p*<0.05 for all comparisons). While, at the 65° shoulder angle, the upper trapezius CoV of the normalized RMS amplitude maps was significantly higher during extension compared to flexion (*p*=0.030).

### 3.2 Motor unit discharge behaviour

**Figures 5** through **8** illustrate, for the different shoulder actions at the 30° shoulder angle, the neural features at the motor unit level that can be extracted using the proposed novel framework. For each muscle, representative MUAP amplitude maps and the corresponding motor unit instantaneous discharge rate are presented. During abduction (**Figure 5**), motor units could be identified only in the deltoid and upper trapezius for this representative participant, with 30 and 19 units decomposed, respectively. For the deltoid, the motor units were distributed across different regions of *array 2*, including anterior (MUAP 1 and MUAP 30), lateral (MUAP 3 and MUAP 5), and posterior regions (MUAP 2 and MUAP 4). In contrast, most of the motor units identified in the upper trapezius were located cranially within the array. Additionally, as seen in the right panel of **Figure 5**, most of the motor units identified from both muscles were tonically active throughout the plateau phase of the action.

**Figure 5:**
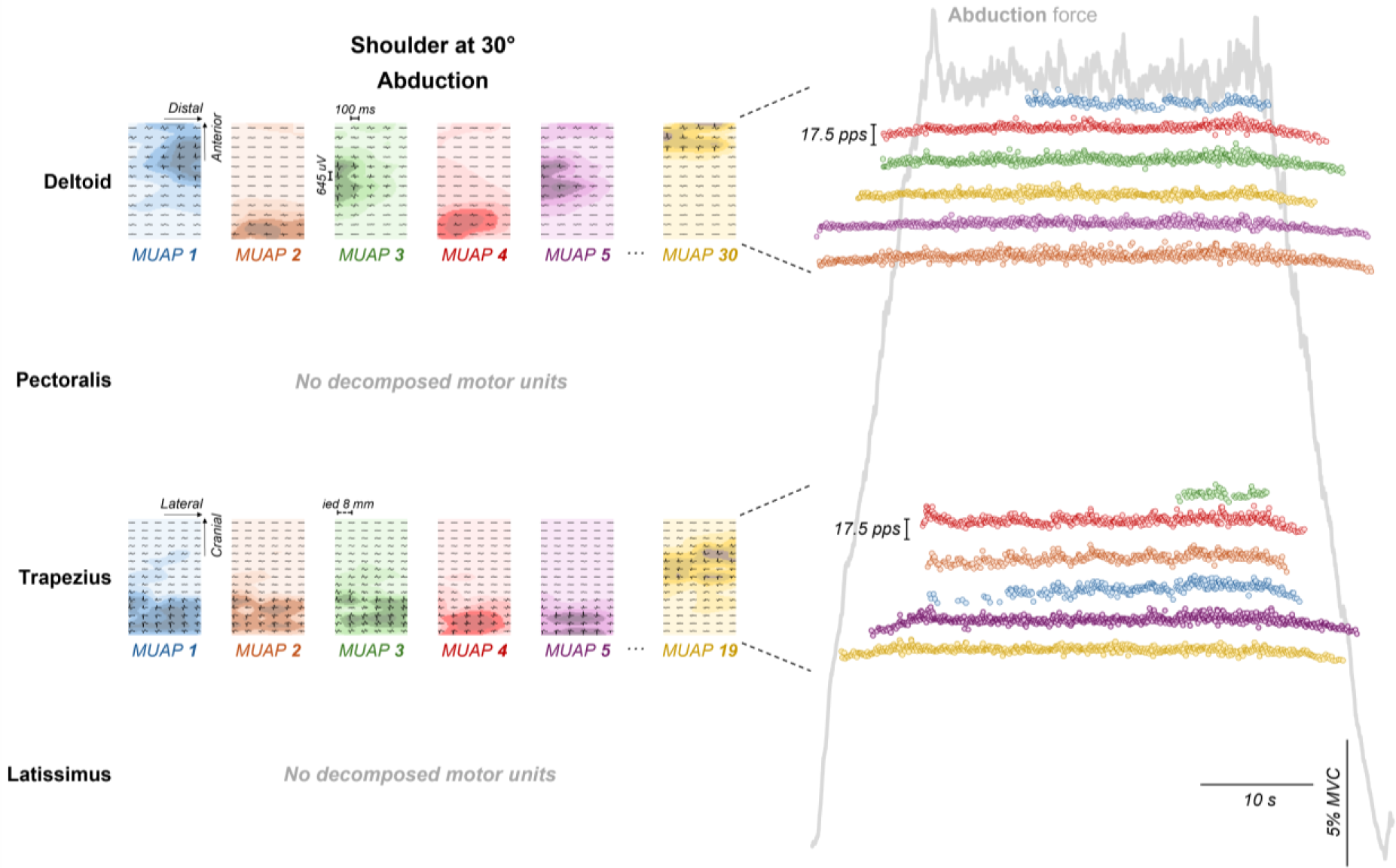
Motor unit action potential amplitude maps and motor unit discharge rate during shoulder abduction at 30°. Motor unit action potentials were obtained using the spike-triggered averaging technique from array 2. For this representative subject, motor units were identified in the deltoid (30 units) and upper trapezius (19 units). No motor units were decomposed from the clavicular pectoralis major or latissimus dorsi (left panel). Abduction force profile (grey line), and motor unit instantaneous discharge rate are shown in the right panel. Separately per muscle, each motor unit is represented with a colour, used both for the MUAP distribution maps and the instantaneous discharge rate.

**Figure 6** is the same representative participant during the adduction action at a 30° shoulder angle. In this condition, motor units were identified in the deltoid (5 units), clavicular pectoralis major (16 units), upper trapezius (2 units), and the latissimus dorsi (13 units). The identified motor units in the deltoid were located across all regions of the deltoid. For the upper trapezius the identified units were located caudally and, in the latissimus, dorsi, as seen in the deltoid, the units were spread across the entire region. Interestingly, the units identified in the clavicular pectoralis major covered the entire *array*, however the majority of the activity following the clavicular pectoralis major line of action. While most of the clavicular pectoralis major and latissimus dorsi motor units were tonically active throughout the plateau phase, some deltoid and two motor units identified in the upper trapezius exhibited more phasic activity.

**Figure 6:**
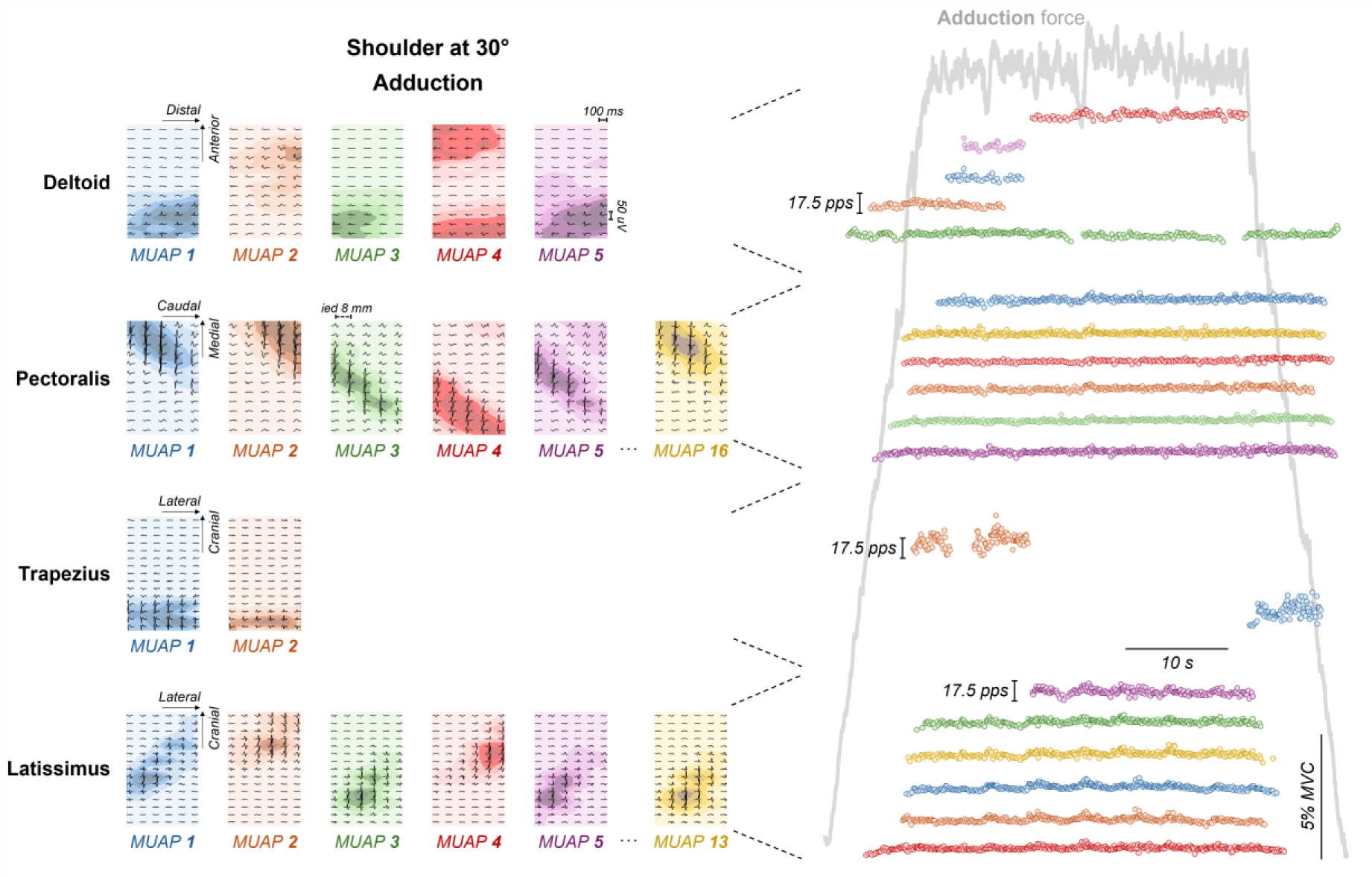
Motor unit action potential amplitude maps and motor unit discharge rate during shoulder adduction at 30°. Motor unit action potentials were obtained using the spike-triggered averaging technique from array 2. For this representative subject, motor units were identified in the deltoid (5 units), clavicular pectoralis major (16 units), upper trapezius (2 units) and latissimus dorsi (13 units) (left panel). Adduction force profile (grey line), and motor unit instantaneous discharge rate are shown in the right panel. Separately per muscle, each motor unit is represented with a colour, used both for the MUAP distribution maps and the instantaneous discharge rate.

**Figure 7** shows the same representative participant during the flexion action at a 30° shoulder angle. In this condition, motor units were identified in the deltoid (20 units), clavicular pectoralis major (21 units), and upper trapezius (16 units), whereas again no motor units were identified in the latissimus dorsi. The identified motor units in the deltoid were predominantly located in the anterior region of the array (e.g., MUAP 1, MUAP 2, and MUAP 5). For both the upper trapezius and clavicular pectoralis major, motor units were distributed across different regions of their respective arrays. While most deltoid and clavicular pectoralis major motor units were tonically active throughout the plateau phase, some upper trapezius units exhibited more phasic activity.

**Figure 7:**
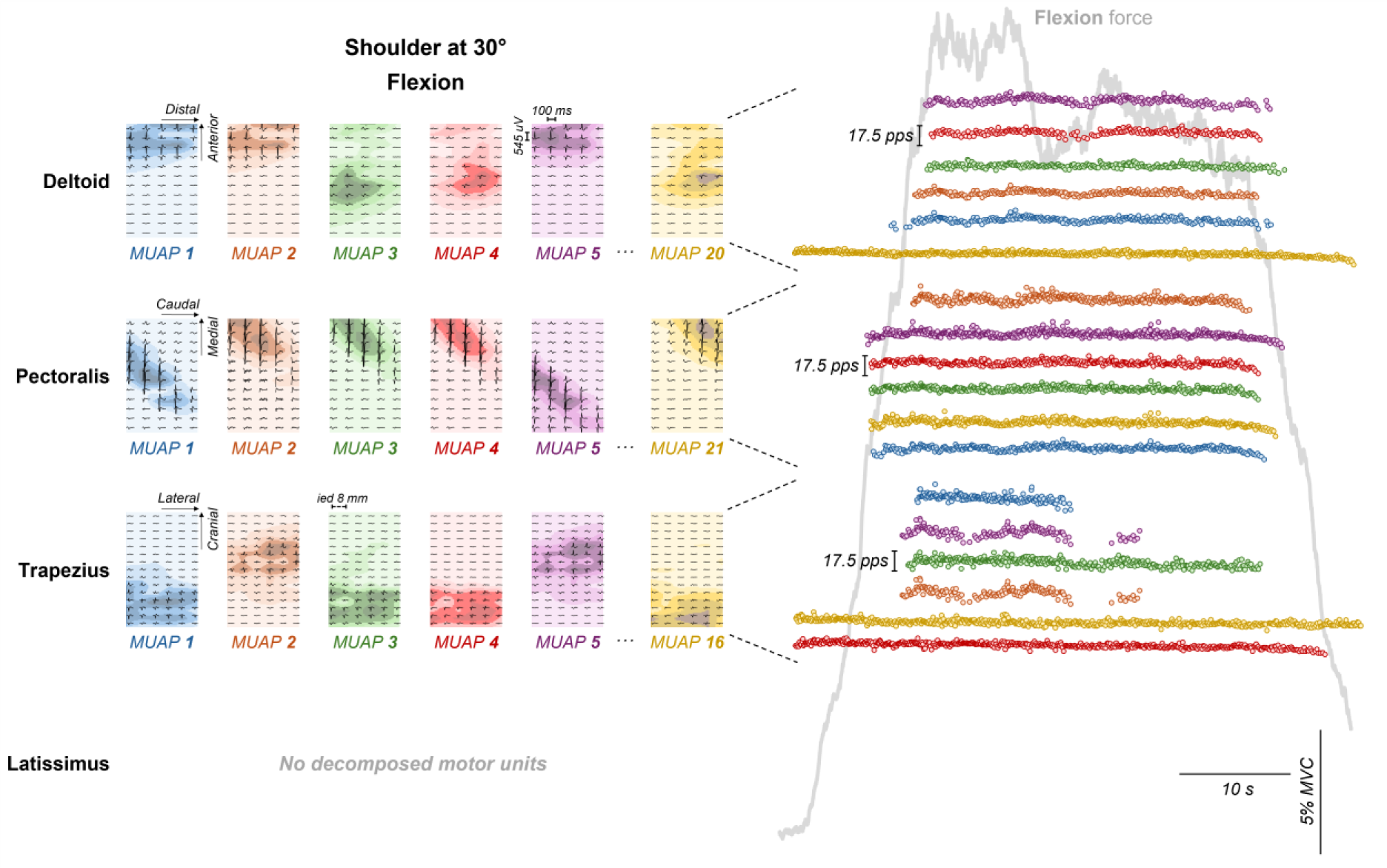
Motor unit action potential amplitude maps and motor unit discharge rate during shoulder flexion at 30°. Motor unit action potentials were obtained using the spike-triggered averaging technique from array 2. For this representative subject (same as in Figure 5), motor units were decomposed from the deltoid (20 units), clavicular pectoralis major (21 units), and upper trapezius (16 units). No motor units were decomposed from the latissimus dorsi (left panel). The flexion force profile (grey line), and motor unit instantaneous discharge rate are shown in the right panel. Separately per muscle, each motor unit is represented with a colour, used both for the MUAP distribution maps and the instantaneous discharge rate.

**Figure 8** illustrates the same representative participant during the extension action at a 30° shoulder angle. In this condition, motor units were identified in the deltoid (9 units), upper trapezius (10 units), and latissimus dorsi (12 units), whereas no motor units were identified in the clavicular pectoralis major. The identified motor units in the deltoid were predominantly located in the posterior region of the array (e.g., MUAP 2, MUAP 3, and MUAP 9). For the upper trapezius, motor unit 1 was dispersed across the entire array, where the other motor units were located more caudally. In the latissimus dorsi, the motor units were distributed across different regions of the array with the majority located more cranially (e.g., MUAP 2, MUAP 4, MUAP 5, and MUAP 12). While most of the deltoid and upper trapezius motor units were tonically active throughout the plateau phase, the latissimus dorsi units exhibited both tonic and phasic activity.

**Figure 8:**
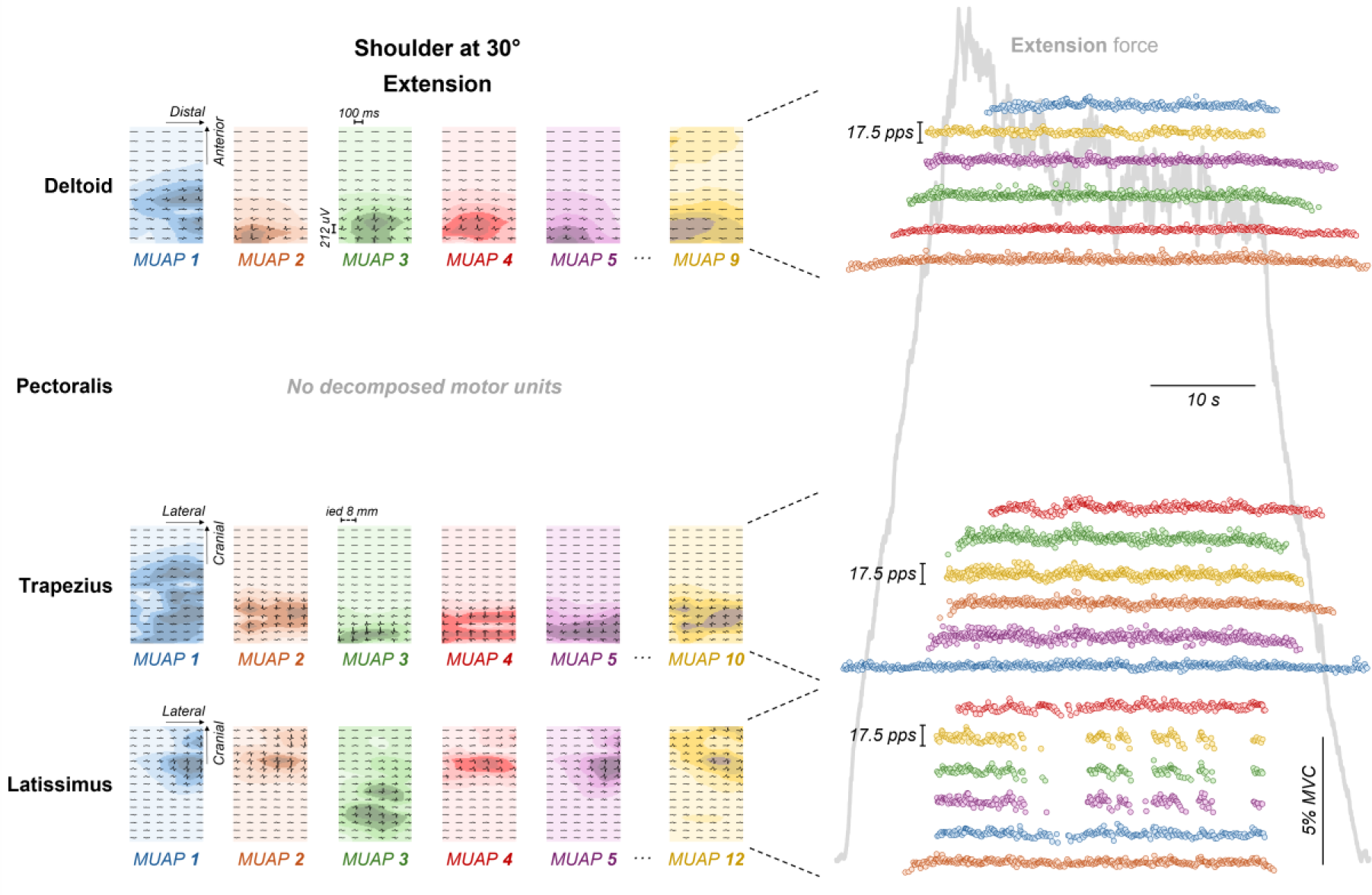
Motor unit action potential amplitude maps and motor unit discharge rate during shoulder extension at 30°. Motor unit action potentials were obtained using the spike-triggered averaging technique from array 2. For this representative subject, motor units were identified in the deltoid (9 units), upper trapezius (10 units) and latissimus dorsi (12 units). No motor units were decomposed from the clavicular pectoralis major (left panel). Extension force profile (grey line), and motor unit instantaneous discharge rate are shown in the right panel. Separately per muscle, each motor unit is represented with a colour, used both for the MUAP distribution maps and the instantaneous discharge rate.

**Table 1** summarizes the motor unit yield across all muscles, actions, and shoulder angles. Overall, a greater number of motor units were decomposed at the 30° shoulder angle compared with 65°. Consistent with the representative case, no motor units were identified in the clavicular pectoralis major and latissimus dorsi during abduction, or in the latissimus dorsi during flexion. In addition, no units were decomposed from the clavicular pectoralis major during extension.

**Table 1:**
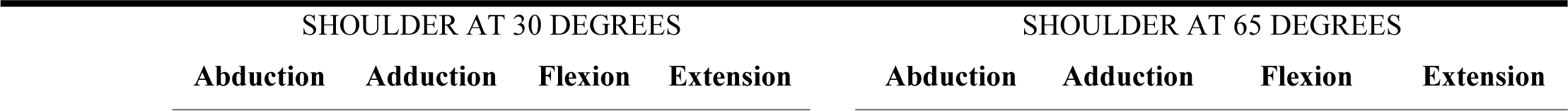
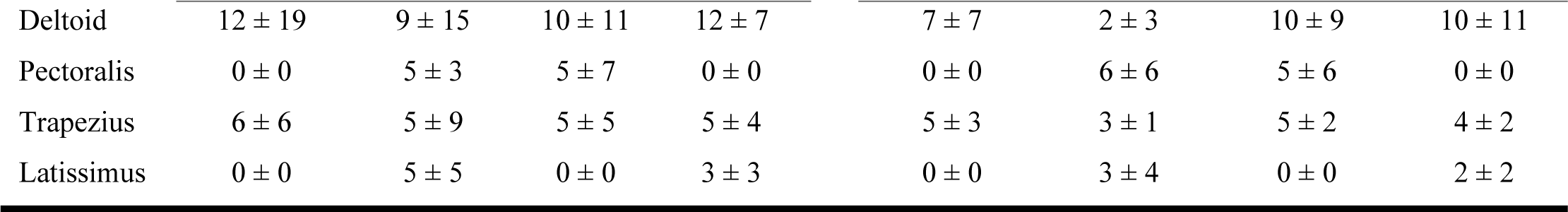
Average ± standard deviation motor unit yield across all muscles (deltoid, clavicular pectoralis major, upper trapezius and latissimus), actions (abduction, adduction, flexion and extension) and shoulder angles (30° or 65°).

**Figure 9** illustrates the group results for the mean motor unit discharge rate for the 30° (**Figure 9a**) and 65° (**Figure 9b**) shoulder angles. For both 30° and 65° there was a statistically significant interaction between muscle and action (LMM, F=3.190, *p*=0.008; LMM, F=6.821, *p*<0.001). The following pairwise comparison were common for both angles. For abduction, there were no significant differences between the mean motor unit discharge rate of the deltoid and upper trapezius (*p*>0.05). During flexion, at the 30° shoulder angle, the motor units of the clavicular pectoralis major had significantly higher mean discharge rate compared to the motor units of both upper trapezius and deltoid (*p*<0.05 for both comparisons). However, during flexion, at the 65° shoulder angle, both the deltoid and clavicular pectoralis major had significantly greater motor unit discharge rates compared to the upper trapezius (*p*<0.001 for both comparisons). For extension, both the deltoid and upper trapezius had significantly higher motor unit discharge rate compared to the latissimus dorsi (*p*<0.05 for both comparisons). When comparing the same muscle across actions at the 30° and 65° shoulder angles, the deltoid had a greater mean motor unit discharge rate during abduction, flexion and extension actions compared to adduction (*p*<0.001 for all comparisons). The mean motor unit discharge rate of the clavicular pectoralis major was higher during flexion than adduction (*p*<0.001), however, the upper trapezius mean motor unit discharge rate during abduction was greater compared to adduction and flexion (*p*<0.05 for both comparisons).

**Figure 9:**
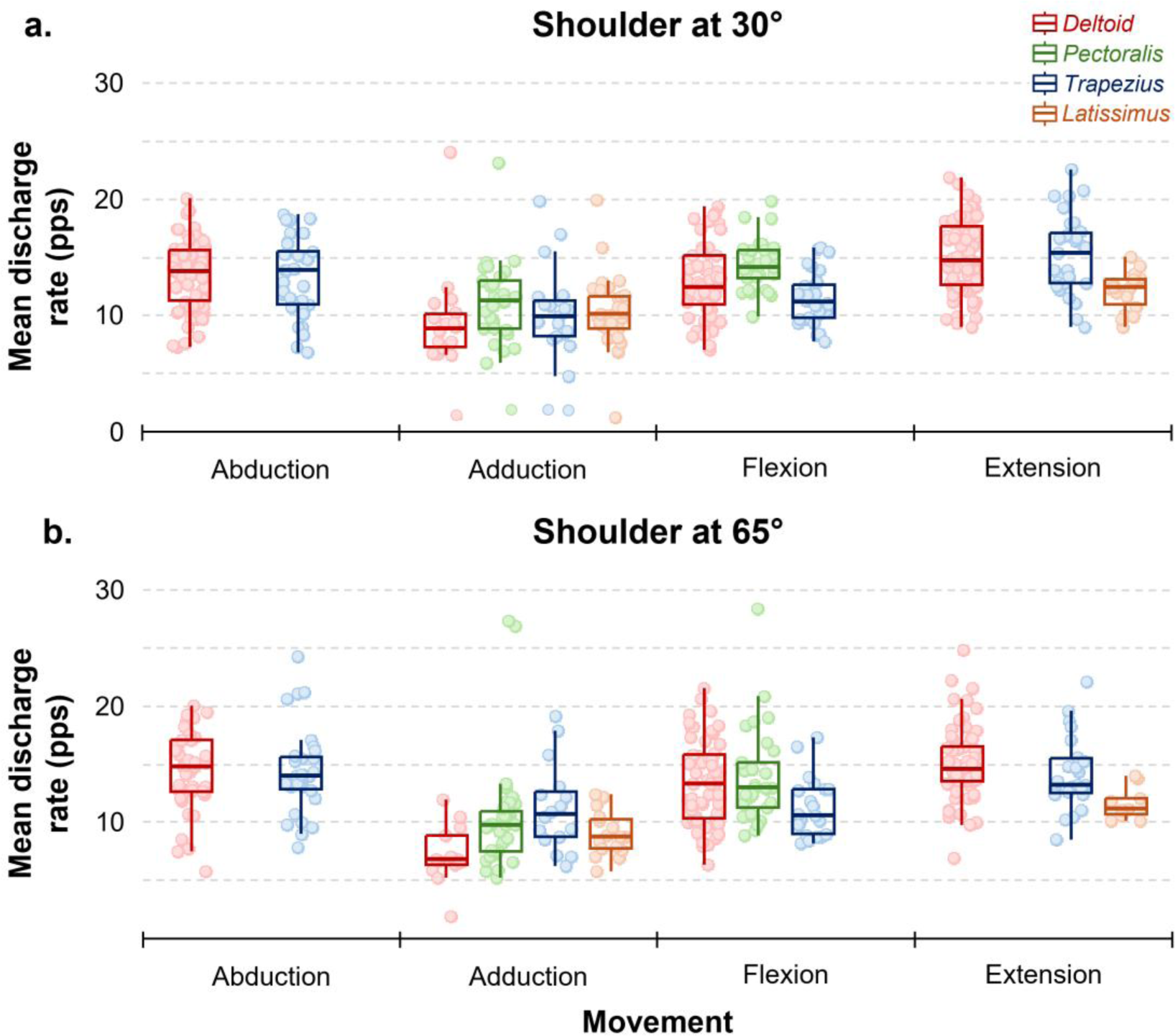
Motor unit discharge behaviour across actions and shoulder angles. Group results for the mean motor unit discharge rate (a and b) during abduction, adduction, flexion, and extension at both 30° (top) and 65° (bottom) shoulder angles. Horizontal traces, boxes, and whiskers represent the median value, interquartile interval, and distribution range, respectively; individual circles correspond to single motor units.

## 4.0 DISCUSSION

The aim of this methodological paper was to propose a novel framework based on the investigation of four isometric shoulder actions at two angles to fully assess the neural control of the superficial shoulder muscles (deltoid, clavicular pectoralis major, upper trapezius, and latissimus dorsi). This framework integrates information across multiple levels of analysis, including global HDsEMG amplitude (normalized RMS), spatial distribution of activation (CoV of RMS amplitude maps), and motor unit discharge behaviour (discharge statistics). To our knowledge, this is the first paper to report motor unit behaviour obtained with non-invasive HDsEMG from the deltoid and latissimus dorsi, and to concurrently assess motor unit activity across all four superficial shoulder muscles during controlled isometric actions. By combining a six-degree-of-freedom load cell, a robotic arm, and evolving HDsEMG technology with motor unit decomposition, the proposed framework demonstrates the feasibility of assessing neural control of the shoulder musculature beyond traditional bipolar EMG approaches. Importantly, this framework provides several examples of the type of extractable neural information that can be obtained when global, spatial, and motor unit analyses are integrated, thereby offering methodological guidelines for future investigations and contributing to the advancement of shoulder neuromechanical and musculoskeletal modelling.

### 4.1 Differences across muscles within the same action

The application of the proposed framework can be integrated into assessments from many disciplines, including but not limited to, sport, physiotherapy and assessment of clinical populations for exercise prescription and rehabilitation. The application of the framework in clinical population, such as stroke and breast cancer survivors, has the ability to assist in monitoring the progression of treatment or the potential negative effects of treatment (Leonardis *et al*., 2022). The proposed framework can be used in addition to assessments with standard bipolar recordings, which has been shown to mischaracterize pectoralis major activity in commonly performed tasks (Lulic-Kuryllo *et al*., 2021*a*, 2023). Therefore, the additional of HDsEMG recordings can provide critical information during assessments.

General consensus is that during complex shoulder actions, most of the muscles will stabilize while others will produce the necessary muscle forces to initiate action (Schenkman & Rugo de Cartaya, 1987; Kelly *et al*., 2002). During shoulder abduction, both the deltoid and upper trapezius exhibited significantly greater global RMS amplitudes compared to the clavicular pectoralis major and latissimus dorsi, consistently across shoulder angles (**Figures 4c, 4d**). This pattern aligns with the primary mechanical roles of the deltoid in humeral elevation and the upper trapezius in scapular stabilization during force generation (Paine & Voight, 1993; Trudeau *et al*., 2022). From a global EMG perspective, these two muscles would therefore be interpreted as the dominant contributors to the abduction action. However, the additional spatial and motor unit analyses revealed important nuances not captured by global amplitude alone. Motor units identified in the upper trapezius were predominantly localized in the caudal portion of the array (**Figures 5 through 8**), indicating a more spatially confined activation strategy (Falla & Farina, 2008*b*). In contrast, the deltoid showed region-specific activation, with motor units distributed across anterior, lateral, and posterior regions of the array, reflecting the engagement of multiple functional compartments within the muscle (**Figures 5 through 8**). Despite these distinct spatial recruitment patterns, the mean motor unit discharge rates did not differ between the deltoid and upper trapezius (**Figures 5 through 8**). This combination of findings suggests that, during abduction, differences between these muscles might not be primarily driven by the magnitude of neural drive, but rather by how that drive is spatially distributed within each muscle. These observations illustrate how the proposed framework allows differentiation of muscle-specific control strategies within a single functional action

During shoulder extension, the deltoid and upper trapezius again exhibited the greatest global EMG amplitude, but this was accompanied by a marked increase in spatial heterogeneity of activation in the upper trapezius, as reflected by a higher coefficient of variation of the RMS amplitude maps (**Figure 4**). This region-specific activation of upper trapezius is in line with previous studies showing higher caudal activation during shoulder actions (Falla & Farina, 2008*a*, 2008*b*). While global measures alone would suggest a secondary role of the upper trapezius in this action, the spatial analysis indicates a more complex activation pattern, potentially reflecting action-dependent stabilization or synergistic control demands (Paine & Voight, 1993; Trudeau *et al*., 2022). At the motor unit level, both the deltoid and upper trapezius displayed higher discharge rates than the latissimus dorsi, despite no differences in normalized global RMS amplitude between them (**Figures 4c, 4d, 9**). This dichotomy between amplitude and discharge behaviour highlights the added value of motor unit analysis for interpreting muscle contributions during multi-joint shoulder actions.

Collectively, these findings highlight the limitation of relying solely on RMS amplitude from bipolar EMG, which, due to waveform cancellation or cross-talk, may not fully capture underlying neural activity (De Luca & Merletti, 1988; Day & Hulliger, 2001; Dimitrova *et al*., 2002; Farina *et al*., 2002, 2010; Keenan *et al*., 2005). Therefore, the use of global EMG measures allows for a general understanding of the duration and magnitude a muscle of interest is activated, but this cannot necessarily indicate the functional role the muscle is contributing to a specific muscle action (Boettcher *et al*., 2010). Therefore, the knowledge of motor unit behaviour during a shoulder action and the region of greatest activation may be essential to provide the information necessary to understand the functional roles of the involved muscles.

### 4.2 Differences within the same muscle across actions

The proposed framework allows the assessment of not only the muscles activity across actions, which we suggest is highly relevant for clinical populations, but also the assessment of the differing activation of individual muscles across actions. This type of assessment is highly relevant for basic research questions regarding the neural control of shoulder actions. To date there is a significant knowledge gap in the neural control of the shoulder, specifically concerning the deltoid and latissimus dorsi (Inglis *et al*., 2025). As was highlighted in the systematic review and meta-analysis by Inglis, Cabral and colleagues (2025), there is much work still to be done to understand the full complexities of the shoulder. Therefore, this proposed framework would be utilized to investigate these complexities, specifically in the deltoid and latissimus dorsi which both have a paucity of research, specifically involving motor unit behaviour non-invasively. The precision of the proposed framework would also allow the tracking of minor activation changes across shoulder angles. These activation adjustments may result from trivial changes in muscle architecture (e.g. pennation angle, muscle length) (Peterson & Rayan, 2011) and moment arm length (Otis *et al*., 1994; Ackland *et al*., 2008) that have significant implications neuromechanically.

When examining the same muscle across different actions, clear action-dependent modulation emerged at both the global and motor unit levels. The deltoid demonstrated significant action-dependent modulation across the investigated shoulder actions (**Figures 4 through 9)**. Global RMS amplitude and mean motor unit discharge rate were significantly greater during abduction, flexion, and extension compared to adduction, confirming its multifunctional role as a primary mover across several directions (Wickham & Brown, 1998). Beyond changes in activation magnitude, MUAP distribution maps for a representative participant revealed a redistribution of activity across the deltoid surface (**Figures 7, 8**), with anterior regions predominantly active during flexion and posterior regions during extension. These findings suggest that neural control of the deltoid involves not only scaling of neural drive but also selective recruitment of different motor unit populations, supporting the notion of functional compartmentalization within large shoulder muscles.

The upper trapezius also exhibited marked action-dependent modulation. Global RMS amplitude was greatest during abduction, consistent with its established role in scapular control (Paine & Voight, 1993; Trudeau *et al*., 2022). However, the spatial distribution of activation varied markedly across actions, with extension eliciting a significantly higher coefficient of variation compared to other actions (**Figures 4e, 4f**). At the motor unit level, discharge rates were highest during abduction and lower during flexion and adduction, suggesting an action-dependent adjustment of the neural drive. Together, these results demonstrate that spatial distribution and motor unit variables provide complementary insights into the neural strategies underlying the upper trapezius activation that are not evident from global amplitude measures alone.

For the clavicular pectoralis major, greater global RMS amplitude and motor unit discharge rates were primarily observed during flexion, whereas little activity was detected during extension (**Figures 4a, 4b**). Similarly, motor units from the latissimus dorsi were identified only during adduction and extension (**Table 1**). Together, these findings demonstrate that muscles contributing to multiple shoulder actions are modulated not only through adjustments in activation level but also through action-specific recruitment and rate coding strategies.

Although motor units were not tracked across actions, the differences in sets of active units between actions strongly suggest action- and angle-dependent recruitment and de-recruitment, consistent with previous findings in other upper-limb muscles (ter Haar Romeny *et al*., 1984). These observations align with previous EMG amplitude evidence showing localized activation of deltoid muscle regions dependent on action direction (Wickham & Brown, 1998, 2012; Brown *et al*., 2007), as well as similar findings in the rotator cuff muscles (Alenabi *et al*., 2018; Lulic-Kuryllo *et al*., 2020). Importantly, the present framework extends these observations by providing direct evidence at the motor unit level.

### 4.3 Methodological implications and future applications

The capability of the proposed framework to map both spatial activation patterns (**Figures 4a, 4b**) and motor unit territories and discharge patterns (**Figures 5** through **8**) provides a powerful tool for probing the neural strategies underlying complex shoulder control. The identification of region-specific motor unit recruitment and tonic versus phasic discharge statistics offers insight into how muscles contribute to both force generation and stabilization during different actions. This is evident during the extension action where the posterior deltoid and upper trapezius are tonically active as they are involved in the action and the latissimus dorsi is both tonically and phasically active as some motor units are recruited to support the action and others (MUAPs 2-4) likely act as stabilizers (**Figure 6**). This multilevel information is particularly valuable for large, multifunctional muscles such as the deltoid and upper trapezius, whose roles cannot be fully understood using global EMG amplitude alone.

Current shoulder musculoskeletal models are based on simplified mechanical analysis and often treat muscles as single force-generating elements (Hughes & An, 1997). The present findings suggest that future modelling studies of the deltoid should consider within-muscle heterogeneity, including compartmentalized segments (compartmentalized within segments: anterior, lateral, posterior) and how this may change based on muscle architecture, moment arm length and action (Hughes & An, 1997). Incorporating motor unit-level information into such models may substantially improve their physiological realism and clinical relevance.

Finally, the differences observed between normalized RMS amplitude and motor unit discharge behaviour at the group level further highlights the limitations of classic EMG-based approaches. Discrepancies between global amplitude and discharge rate plausibly reflect factors such as amplitude cancellation, the limited spatial coverage of HDsEMG arrays, and potential cross-talk from adjacent muscles (De Luca & Merletti, 1988; Day & Hulliger, 2001; Dimitrova *et al*., 2002; Farina *et al*., 2002; Keenan *et al*., 2005). By directly assessing motor neuron output, the proposed framework provides a more precise assessment of neural drive and regional muscle involvement. This methodological advance has important implications for both basic research and clinical applications, particularly in populations where subtle alterations in neural control may underlie functional impairment and where targeted rehabilitation strategies are required.

### 5.0 PRACTICAL IMPLICATIONS

- Using the proposed framework, rehabilitation programs can be designed based on the specific actions that isolate the motor unit activity, not the global muscle activity, to target specific impairments better
- The proposed framework provides a roadmap for stroke rehabilitation, where the focus may need to be centred more on motor unit recruitment and rate coding rather than on general activity within a portion of the muscle.
- The proposed framework provides a detailed description of the equipment used during the data collection including the testing jig, recording equipment, array locations and actions at each joint angle to be reproduceable for scientists and clinicians investigating recruitment of the shoulder musculature.
- The use of the robotic arm in the proposed framework allows for the testing jig and load cell to be situated so the position of the humerus, glenohumeral joint angle, muscle and moment arm length do not change across actions, only the jig moves to allow the recording of the forces produced during the specified action direction.

## 6.0 LIMITATIONS

Despite the rigor taken to control the specific shoulder actions and position, several limitations should be acknowledged. First, although participants were secured with straps and a rigid jig, the inherent flexibility of the shoulder complex may have introduced small variations in action execution. While this may reduce experimental control, it also increases ecological validity by better replicating strategies used in daily activities. Second, only two joint angles within the mid-range were tested, limiting extrapolation to positions closer to 0° or 180°. Third, contractions were restricted to 30% MVC, chosen to reflect moderate force demands of activities of daily living, but additional work is needed across lower and higher force levels. Fourth, the sample included only one female participant, precluding sex comparisons. Finally, only four major superficial shoulder muscles were recorded. The inability to assess, particularly the scapular stabilizers and rotator cuff, is a key omission given their likely contribution during extension and adduction actions.

## 7.0 CONCLUSIONS

The present study demonstrates that it is feasible to record and analyse motor unit behaviour of major superficial shoulder muscles using HDsEMG during controlled isometric actions. The framework combines motor unit analyses with precise force measurements to provide complementary insights into neural control strategies. These methodological advances are essential for the development of comprehensive biomechanical models of the shoulder that incorporate not only anatomical and mechanical properties but also neural control strategies.

## AUTHOR CONTRIBUTIONS

- Conceptualization: JGI, SR, HVC, TLK, CRD, FN
- Data curation: JGI, SR, HVC
- Formal analysis: JGI, SR, HVC
- Funding acquisition: JGI, FN
- Investigation: JGI, SR, CC
- Methodology: JGI, SR, HVC, RP, TLK, CRD, FN
- Project administration: FN
- Writing – original draft: JGI, SR, HVC, FN
- Writing – review and editing: JGI, SR, HVC, CC, RP, TLK, CRD, FN

